# Testing the peak-end rule in bumblebees: lack of preference for a higher-reward sequence when the final reward is disappointing

**DOI:** 10.1101/2024.11.25.623196

**Authors:** Mélissa Armand, Pauline Riemer, Sophia Hagn, Jana Schweiger, Tomer J. Czaczkes

## Abstract

The peak-end rule describes the tendency to evaluate experiences by their most intense and final moments, rather than considering the entire experience as a whole. While this cognitive bias is well-established in humans, studies on nonhuman animals are very limited. Bumblebees make foraging decisions largely based on past experiences, but whether peak-end effects influence their subsequent flower choices is still unknown. Here, we trained individual *Bombus terrestris* workers on two artificial flower types, alternating blue and yellow, over 12 consecutive foraging bouts. One flower type offered a sequence of three high-quality rewards (25 μL drops of 50% w/w sucrose solution: “consistent” sequence), while the other provided the same sequence but ended with an additional, lower-quality reward (25 μL drop of 20% w/w sucrose solution: “poor end” sequence). We then tested the bees’ flower type preference in a final binary choice. Bees showed a strong preference for blue flowers, both in their initial and overall visits. Across all visits during a 1-minute period, they also favoured flowers associated with the “consistent” sequence, though this preference was significant only when these flowers were yellow. Interestingly, despite offering more sucrose per foraging bout, bees did not favour the “poor end” sequence flower. This study is, to our knowledge, the first to investigate peak-end effects in an insect. How bees evaluate sequential rewards when foraging remains largely unexplored, yet could provide valuable insights into nectar distribution and plant-pollinator co-evolution.

## INTRODUCTION

Animals encounter countless options to choose from throughout their life, for example when selecting a nest, partner, or food sources when foraging. Such resources are typically encountered sequentially, requiring individuals to recall and evaluate each option when making decisions. Since memory is a costly and limited resource, individuals may base their judgments on key moments from past experiences rather than the entire experience. Fredrickson and Kahneman (1993) showed that people often rely on the most intense moment (the peak) and the final moment (the end) of an event — a phenomenon known as the peak-end rule. This heuristic is well-established and robustly supported by extensive human studies (see meta-analysis by Alaybek et al., 2022). Specifically, peak-end effects have been shown to strongly influence judgment of sequences, including series of rewards or events (Ross and Simonson, 1991; Ariely, 1998; Chapman, 2000; Lau-Gesk, 2005).

Among studies on the peak-end rule, “end effects” are the most frequently described. People tend to prefer experiences with positive endings (Loewenstein and Prelec, 1993; Diener et al., 2001; Just et al. 2015), or sequences that improve over time (Ariely, 1998; Chapman, 2000; Lau-Gesk, 2005). End effects are likely more prevalent because they are easier to test, whereas identifying peaks, as the most intense moments of an experience, can be more challenging. Additionally, the strong impact of endings on judgment and memory may be attributed to recency effects, arising simply from being experienced last (Ebbinghaus, 1885). Intriguingly, multiple studies have shown that a less favourable end can lower the perceived value of an otherwise superior experience. For instance, in the classic study by Kahneman et al. (1993), participants preferred to repeat a longer cold-water trial with a less painful ending over a shorter, consistently painful one. Similarly, Schreiber and Kahneman (2000) showed that adding a milder final segment to an unpleasant episode led to a more favourable overall memory, even though total discomfort increased. For positive experiences, people rated a happy life ending abruptly as better than a similar one extended with mildly happy years, despite having more total pleasure (Diener et al., 2001). Do et al. (2008) found that adding a lower-quality gift to a high-quality gift reduced satisfaction compared to receiving only the high-quality gift, despite the increase in total value.

However, very few studies have investigated peak-end effects in other animals. Research so far focused solely on primates, with inconsistent findings (Xu et al., 2011; Jung and Kralik, 2013; Blanchard et al., 2014; Egan Brad et al., 2016), but see a recent study on calves (Ede et al., 2023). Choices based on sequential experiences are especially relevant to nectarivores like bumblebees, which forage by moving from flower to flower within an inflorescence or flower patch, experiencing a sequence of rewards during each visit. Although prior experiences influence their foraging decisions (Dukas and Real, 1993; Biernaskie et al., 2009), whether bees are affected by peak-end effects when evaluating rewards and selecting flowers remains unknown. Notably, evidence for end effects in animals is inconsistent. Some studies showed that monkeys instead tend to choose sequences starting with high-value rewards over the reverse, indicating a preference for instant gratification (Xu et al., 2011; Jung and Kralik, 2013). Similarly, Ede et al. (2023) found no peak or end effects in calves’ memory of pain. In contrast, Egan Brad et al. (2016) found that capuchins, like humans, tend to favour sequences ending with pleasurable rewards but do not intentionally structure events to maximize this effect. Blanchard et al. (2014) found that rhesus monkeys preferred sequences with high peak values and were especially sensitive to rewards at the end, to the extent that a small additional reward could decrease overall preference for the sequence. This aligns with previous findings on how a less satisfying end affects overall preference in humans.

The peak-end rule may be of direct ecological importance. Some animals, like bees, encounter sequential food rewards when foraging, requiring them to recall and evaluate flower options based on prior experiences for their subsequent foraging decisions. For instance, bumblebees assess each flower independently and adjust their choices based on reward variability and prior knowledge (Dukas and Real, 1993; Chittka et al., 2003; Biernaskie et al., 2009). Like humans, bees were also shown to be susceptible to recency effects (Prabhu and Cheng, 2008; Nityananda and Chittka, 2021). When foraging, bees rely on various heuristics to simplify decision-making. Bumblebees, for example, may use a simple “nearest-neighbour” rule, moving to the closest available flower (Ohashi et al., 2007; Saleh and Chittka, 2007). Alternatively, they may refine foraging routes through trial-and-error, reinforcing shorter paths and gradually creating efficient routes between flowers (Reynolds et al., 2013). In uncertain conditions, they have been shown to adopt a “copy-when-uncertain” heuristic, relying on social cues over personal information when nectar distribution is unpredictable (Smolla et al., 2016). In a study by MaBouDi et al. (2020), bumblebees trained to differentiate between larger and smaller shapes for a reward used a simple “win-stay/lose-switch” rule-of-thumb, improving their decision-making efficiency. In honeybees, the addition of an inferior option biased bees toward choosing an option similar to it, appearing more attractive by comparison — a heuristic known as the decoy effect (Shafir et al., 2002; Tan et al., 2014). In nature, bees often forage on vertical inflorescences like foxgloves or wild sage, typically beginning at the lowest flowers and moving upward (Pyke, 1979; Waddington and Heinrich, 1979; Best and Bierzychudek, 1982). This behaviour results in a clear sequence of sub-events (individual flowers) per visitation event (inflorescence). Consequently, the overall perception of an inflorescence may be influenced by the order in which flowers with varying rewards are encountered. If the final flower contrasts poorly with the preceding ones, could it lower the perceived value of the entire inflorescence?

Here, we investigated whether bumblebees’ preferences between two flowers were influenced by the quality of the final reward on each flower. Our experiment was inspired by the study of Do et al. (2008), who tested the peak-end rule in humans. In their experiments, participants who received a high-quality gift followed by a lower-quality one reported lower overall satisfaction than those who received only the high-quality gift, despite the second gift objectively increasing the total value. We trained individual *Bombus terrestris* bees on two distinct flowers, each offering a different sequence of nectar rewards: one with three high-quality (50% w/w) drops of sucrose solution, and the other with an additional lower-quality (20% w/w) sucrose solution drop. Although the second flower type offered a greater total amount of sucrose, we hypothesized that bees would prefer the flower associated with the sequence of three high-quality rewards, as it ends better than the sequence with a lower-quality final reward, despite its lower total sucrose value.

## MATERIAL AND METHODS

### Bees

Commercial *Bombus terrestris* colonies purchased from Koppert (The Netherlands) were used for the experiment. The bees were housed in plastic nestboxes (23 × 21 × 12 cm) which consisted of their original transport container, modified with a removable Plexiglas lid for easy handling of individuals. The colonies were kept under controlled laboratory conditions, maintained at 22 – 24°C with a 14:10 light:dark cycle, and with unlimited access to pollen. Each nestbox was connected to a flight arena (60 × 35 × 25 cm) via a transparent tube, leading to a small chamber (6 × 5 × 3 cm) equipped with shutters to regulate the movement of bees in and out of the nestbox. The bee selected for the experiment was allowed through a second tube connecting the chamber to the flight arena (see apparatus in **Fig. 1**). Between experimental sessions, bees had unrestricted access to the arena which contained six artificial flowers offering 20% w/w sucrose solution, replenished regularly throughout the day. Bees that regularly foraged on the flowers were marked on the thorax with uniquely numbered, coloured tags and selected for the experiment. A total of 71 bees were tested: 37 bees from three colonies between November and December 2022, and 34 bees from three additional colonies between April and May 2024 (see Supplement **S1** for further colony details).

**Figure 1.**
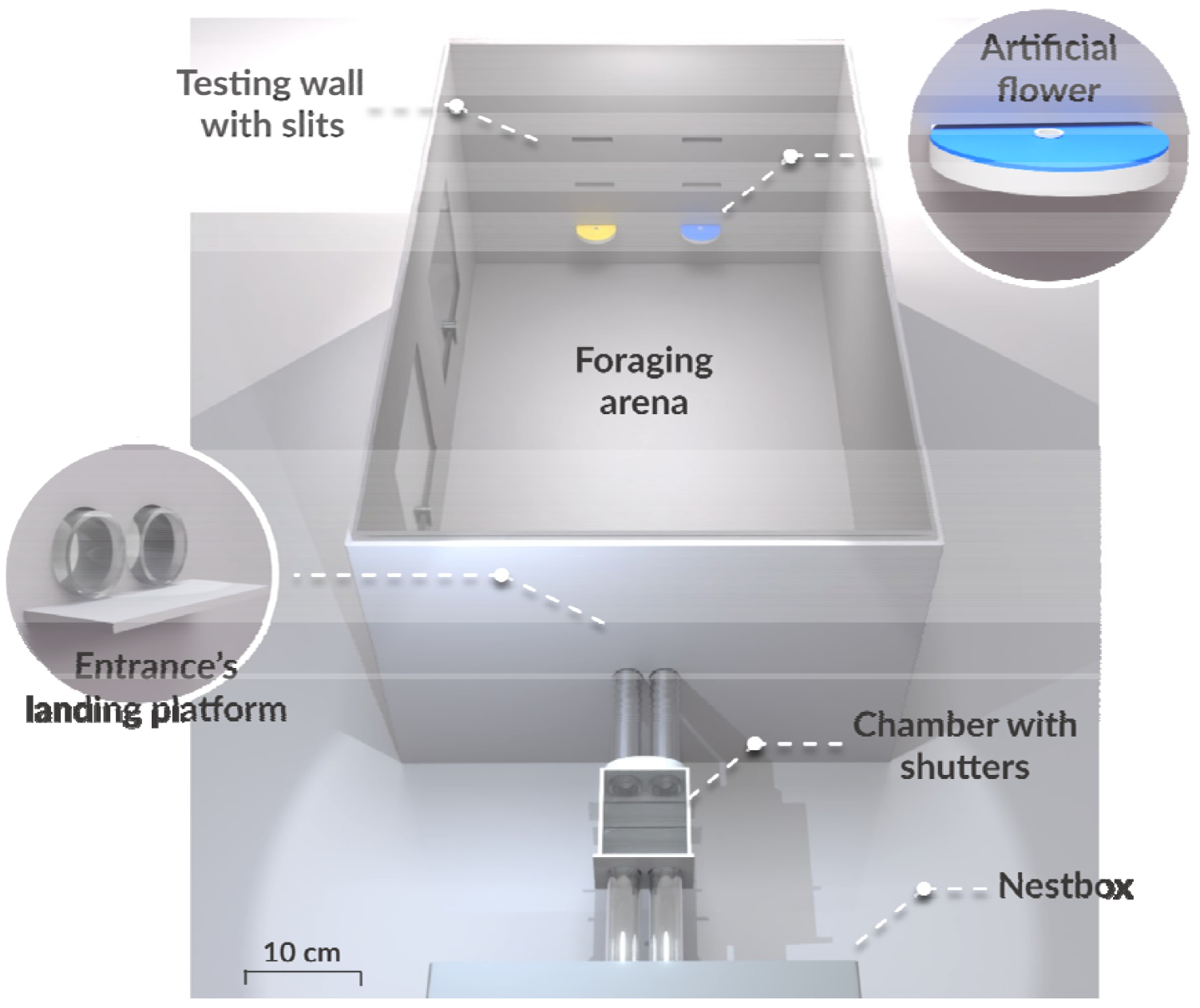
Top view of the scaled experimental setup. The 3D model (created in Blender) illustrates the final binary choice test, with the blue flower positioned on the right side and the yellow flower on the left, based on the bee’s left-right orientation.

### Artificial flowers

Each artificial flower used in experiment consisted of a white plastic disc (diameter: 3.8 cm, thickness: 6 mm) topped with a 2 mm piece of coloured rubber foam. At the centre of each flower, a small opaque white resin cup (diameter: 4 mm, depth: 6 mm) was inserted into a pre-cut hole, and sucrose solution was pipetted into the cup (see **Fig. 1**). The flowers used during the experiment were either blue (with a peak reflectance at approx. 450 nm) or yellow (with a peak reflectance from approx. 540 to 700 nm, see reflectance curves in Supplement **S2**). Between experimental sessions, neutral white flowers were used to prevent the bees from associating experimental colours with sucrose rewards. The flowers were mounted horizontally into slits along the back wall of the flight arena, facing the entrance to ensure they were visible from the bees entering the foraging area. The wall contained a total of six slits arranged in three rows of two, with the lowest row positioned 5 cm above the arena floor and the highest row at 15 cm. Slits within the same row were spaced 6.5 cm apart (see **Fig. 1** below). To collect sucrose solution, the bees were required to fly to the flowers, land, and extract the solution from the resin cups.

### Main experiment

Individual bees completed twelve consecutive foraging bouts, with blue and yellow flowers presented alternately in each bout. Each flower type appeared six times before a final binary choice test, where both flowers were presented simultaneously. The flowers corresponded to two nectar reward sequences:

1. **Consistent sequence**: Three successive flowers (either all blue or all yellow), each offering 25 μL of high-quality (50% w/w) sucrose solution;
2. **Poor end sequence**: Three flowers identical to the consistent sequence, followed by a fourth additional flower offering 25 μL of lower-quality (20% w/w) sucrose solution.

This design ensured that bees experienced rewards sequentially. Each flower provided a total amount of sucrose solution per bout (75 μL for the consistent sequence and 100 μL for the poor end sequence), both below the average crop capacity of bumblebees (120 – 180 μL; Lihoreau et al., 2010), ensuring that all rewards could be collected by the bees.

At the start of each bout, the first flower was positioned in the lowest row of the wall, either in the left or right slit, ensuring its colour was clearly visible to the bee from above when entering the arena. Once the bee began collecting the reward, the second flower in sequence was gently inserted into the opposite slit by an experimenter stationed behind the arena. This process was repeated for the third flower, and for the fourth in the case of the poor end sequence (see video examples of bees foraging in Supplement **S3**). After the bee finished interacting with the final flower, the flower was removed, signalling the end of the bout and encouraging the bee to leave the arena. After each bout, all flowers were replaced with clean ones to prevent scent marks left by bees from affecting subsequent flower choices (Goulson et al., 2000; Saleh et al., 2007).

The positioning of the flowers (left/right slit) remained consistent across all twelve bouts for each bee, with the side order reversed for each flower type. For instance, blue flowers were always presented first on the left, while yellow flowers were presented first on the right. If a bee did not fully collect or engage with the fourth, lower-reward flower in the poor end sequence, it was allowed up to three probing attempts before the flower was removed, ensuring sufficient exposure to the lower-quality rewards. Notably, most bees (60 out of 71) rejected all lower-quality rewards, disengaging from the drops in less than a second (engagement time for a high-quality exceeded five seconds). However, every bee still probed the lower-quality drops in each bout.

After completing all twelve foraging bouts, the bee performed a final bout, where it had to choose between the two flower types placed side by side in the wall, both unrewarded. The flower positions remained consistent with the earlier bouts to help the bee anticipate each flower’s location and reduce the likelihood of rushed or random choices. The first flower the bee landed on was recorded as an indicator of its preference.

### Control experiment

A control experiment was conducted before the main trial to confirm whether the setup allowed bees to associate blue and yellow flowers with rewards. The setup mirrored the main experiment, but one flower type was associated with a sequence of three high-quality rewards (25 μL of 50% w/w sucrose solution), while the other was associated with three lower-quality rewards (25 μL of 20% w/w sucrose solution). This clear difference in reward quality allowed us to test whether bees could correctly identify and preferentially choose first the flower associated with the higher-quality reward sequence in the final test. All tested bees (n = 10) successfully landed first on the flower type associated with high-quality rewards during the binary choice test.

### Data analysis

Each foraging bout was video-recorded using a Sony HDR-CX220 camera positioned above the flight arena. Bee behaviour was analysed using the event-logging software BORIS (v8.6). For each bee, we recorded: (1) reward collection (continuous licking of a sucrose solution drop for more than five seconds), (2) first flower visit (landing on the flower surface) in the binary choice test, and (3) all flower visits over a 1-minute period during the binary choice test.

Data processing was performed with Python (v3.11, Python Software Foundation, 2023) using the libraries *pandas* (McKinney, 2010) for data structuring, and *seaborn* (Waskom, 2021) and *Matplotlib* (Hunter, 2007) for data visualisation. Statistical analyses were conducted in R (v4.1, R Core Team 2022) using the *glmmTMB* package (Brooks et al., 2017) for Generalised Linear Mixed Models (GLMM), and *emmeans* (Lenth, 2020) for post-hoc tests. Model residuals were evaluated with the *DHARMa* package (Hartig, 2020). Datasets from both experimental sessions (2022 and 2024) were combined for analysis, with session treated as a random effect to account for variations in experimental conditions. Individual bees, nested within their respective colonies, were also included as random effects to control for intercolony variability. Potential confounding effects to sequence type, such as flower colour (blue or yellow), flower side (left or right slit in the arena), and flower exposure order (presented in the first or second foraging bout), were also tested for any influence on the bees’ flower visits. Complete statistical analyses and datasets are available on Zenodo (https://doi.org/10.5281/zenodo.14068426).

## RESULTS

### 1 Flower colour influenced the bees’ first flower choice

We first conducted a preliminary analysis to assess the influence of confounding flower factors on the bees’ first visit during the binary choice test. Specifically, we examined whether the proportions of first visits differed by flower colour (blue or yellow), flower side (left or right), and exposure order (first or second). Among these factors, only flower colour showed significant differences in proportions, with bees visiting blue flowers more frequently than yellow flowers (binomial test: probability of visiting blue = 0.77, *p* < 0.0001, N = 71 visits).

We then tested whether the sequence type (consistent or poor end) influenced the bees’ first flower choice using a generalized linear mixed model (GLMM), accounting for flower colour as a confounding factor and including random effects for colony and experimental session. The sequence type did not significantly affect the bees’ first choice, regardless of the associated flower colour (GLMM, binomial family; yellow flowers: Intercept = 0.51, SE = 0.52, *z* = 0.99, *p* = 0.32; blue flowers: Intercept = 0.04, SE = 0.27, *z* = 0.13, *p* = 0.89, N = 71 visits). When the consistent sequence flower was yellow, 62.5% of bees chose it first, compared to 50.9% when it was blue (**Fig. 2A**). Although bees showed a higher tendency to select the consistent sequence flower when it was yellow, this difference was not statistically significant (post hoc Tukey test: odds ratio = 1.61, SE = 0.936, *z* = 0.814, *p* = 0.415, N = 71 visits).

**Figure 2.**
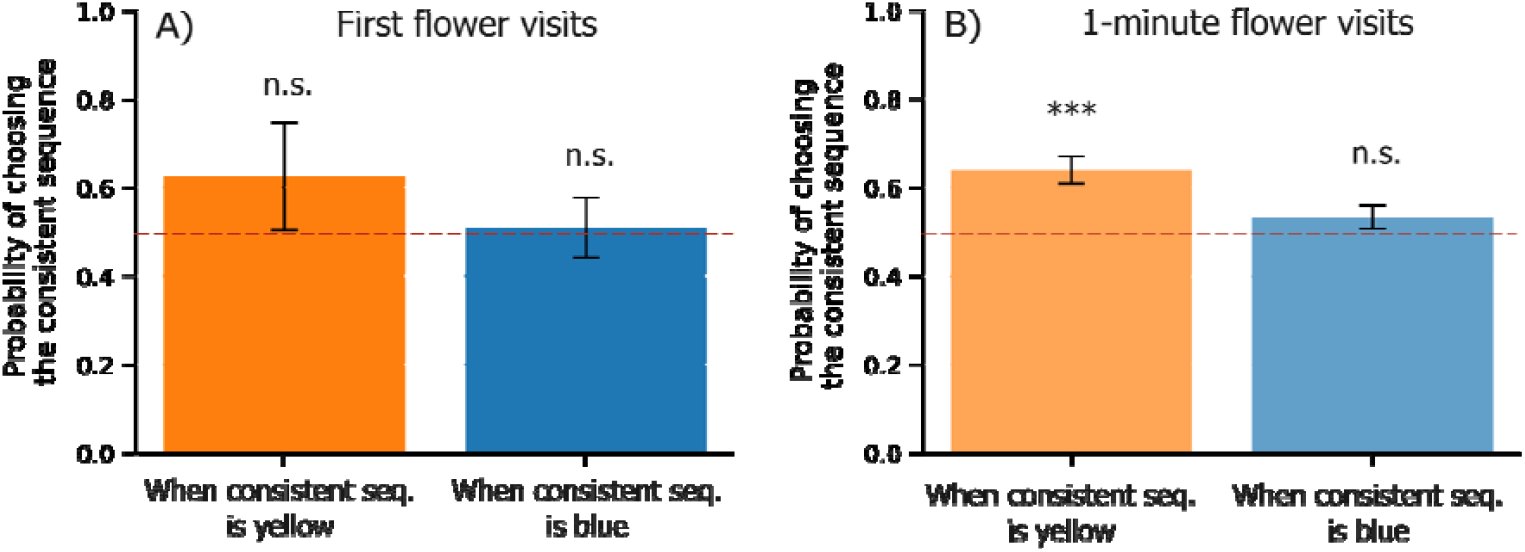
Predicted probabilities of visiting the consistent sequence flower for each flower colour. **A:** Proportion of choices in the first flower visit, when the consistent sequence flowers are blue or yellow (N = 71 visits); **B:** Proportion of choices across all flower visits over a 1-minute period, when the consistent sequence flowers are blue or yellow (N = 605 visits). Error bars indicate the standard errors of the predicted probabilities, and statistical significance is indicated as: **\*\*\*** (*p* < 0.001), and **n.s**. (*p* > 0.05).

### 2 The interaction between flower colour and sequence type influenced bees’ overall flower visits

Next, we considered all flower visits made by the bees over a 1-minute period in the binary choice test. We examined the proportion of overall visits using the same factors as in 1). Only flower colour significantly affected choice proportions, with bees visiting blue flowers more frequently than yellow flowers (binomial test: probability of visiting blue = 0.60, *p* < 0.0001, N = 605 visits).

We then tested the effect of sequence type while accounting for flower colour and random effects of colony and experimental session. The bees visited significantly more flowers associated with the consistent sequence when the flower was yellow (GLMM, binomial family: Intercept = 0.57, SE = 0.13, z = 4.26, *p* < 0.0001, N = 605 visits), but not when the flower was blue (Intercept = 0.13, SE = 0.11, z = 1.26, *p* = 0.209, N = 605 visits). When the consistent sequence was associated with yellow flowers, it accounted for 63.9% of visits, compared to 53.3% when associated with blue flowers (**Fig. 2B**). This difference was significant (post hoc Tukey test: odds ratio = 0.645, SE = 0.11, z = -2.577, *p* = 0.010, N = 605 visits).

## DISCUSSION

This study examined whether bumblebees’ preference between two flowers was influenced by the quality of the final reward received during a foraging bout on each flower type, assessing their susceptibility to order effects such as the peak-end effect. Contrary to our hypothesis, bees did not show a clear preference for the “consistent” flower with three high-quality rewards over the “poor end” flower with an additional, lower-quality reward at the end. Instead, their flower preference was largely driven by colour, with bees favouring blue flowers in both initial and overall visits during the binary choice test. However, tellingly, bees also did not favour the poor end sequence, despite its higher total value. We also note that, when considering all visits, bees showed a preference for flowers associated with the consistent sequence, but this was significant only when the flowers were yellow. Our results align with most findings on nonhuman animals, as studies in monkeys have shown inconsistent endpoint effects in preferences for food sequences (Xu et al., 2011; Blanchard et al., 2014; Egan Brad et al., 2016).

The lack of preference for the flower type with a higher total sucrose solution is intriguing and may reflect an endpoint effect, where a disappointing ending lowers the perceived value of an option. Similarly, in Kahneman et al. (1993), over one-third of participants did not favour the shorter trial with less total pain, nor did they prefer the longer trial with a less painful ending. Such absences of preference for the more profitable option are paradoxical and defy logical expectations. In fact, humans and animals have long been shown to deviate from normative rationality in their decision-making (Tversky and Kahneman, 1989; Shafir and LeBoeuf, 2002; Bateson et al., 2003). For instance, Solvi et al. (2022) demonstrated that bumblebees prioritise the remembered ranking of a flower’s reward over its actual value, opting for flowers they recall as superior even when the rewards are equivalent.

In our experiment, most bees did not collect the lower-quality rewards in the poor-end sequence, even though the total volume of sucrose solution was well below their average crop capacity (Lihoreau et al., 2010). Furthermore, the quality of the “poor” reward (20% w/w sucrose solution) is considered adequate in natural flowers for bees (Pamminger et al., 2019), and we maintained the bees on this same sucrose concentration in the lab. Foraging bees often reject downshifted rewards after experiencing higher-quality ones, a behaviour known as the negative incentive contrast effect (Bitterman, 1976; Waldron et al., 2005; Townsend-Mehler et al., 2011; Hemingway and Muth, 2022). Bumblebees, in particular, have been shown to quickly abandon a previously rewarding food source to seek out new food sources instead of returning to lower-quality options (Townsend-Mehler and Dyer, 2012). This tendency suggests that a negative incentive contrast effect may have reduced the perceived appeal of the additional sugar offered in the poor-end sequence.

Although most bees in our experiment did not fully collect the downshifted flower, each bee still probed the extra reward, thus making them aware of the overall higher absolute value of this flower type, even if they chose not to take advantage of it. As an alternative to an order effect, this lack of preference might reflect an aversion to variability; studies show that bumblebees generally prefer flowers with consistent rewards, even when both options provide the same average reward (Waddington et al., 1981; Harder and Real, 1987). The lower-value final reward may have made the flower type appear unpredictable or “risky”, further discouraging bees from choosing it.

While the exact mechanisms remain unclear, the lack of preference for the consistent sequence over the poor-end one is still informative. Our findings suggest that any peak-end effect, if present, was likely small and largely masked by colour preference. We did find that bees tended to visit flowers associated with the consistent sequence more often, but this was statistically significant only over numerous visits and only when flowers were yellow, when colour preference was likely less influential. Bees have indeed a strong innate preference for blue flowers (Gumbert, 2000; Raine and Chittka, 2007), as reflected in our results. However, such preferences are flexible and can be reversed through learning (Gumbert, 2000). In our control experiment, where flowers had sharply contrasting reward qualities, bees consistently chose the highest-rewarding flower first, overriding their preference for blue.

Although bees showed a preference for consistent sequence flowers in their overall visits, this finding should be considered with caution. Since both flowers were unrewarded in the final binary choice test, bees likely checked the other flower after finding their preferred one empty, and repeated visits may reflect rushed or frustrated behaviour rather than genuine preference. However, similarly, Bortot et al. (2020) analysed honeybee preferences by evaluating total visits made within one minute rather than just the first visit, discovering a strong preference for one flower type that was not apparent in the initial choice. The cumulative choices animals make may thus reveal preferences that their initial choices do not show.

The main challenge in demonstrating peak-end effects in nonhuman animals likely stems from the methodology used to test such biases. In classic experiments, people evaluate experiences retrospectively (*e*.*g*.: Kahneman et al., 1993; Ariely, 1998; Do et al., 2008), where participants are asked to report their overall feelings about various options. This approach is difficult to implement with nonhuman animals, which are typically required to choose their preferred options directly. For example, rhesus monkeys asked to select their favourite food sequence displayed a “peak-first” bias, preferring sequences that offer an immediate high-value reward over those that conclude with the most favourable one (Xu et al., 2011; Jung & Kralik, 2013). Egan Brad et al. (2016) found that human adults, children, and capuchin monkeys preferred sequences that ended with the most pleasurable reward; however, none of these groups intentionally structured events to enhance this end effect. In the study by Diener et al. (2001), where participants rated the desirability of fictional lives that ended either abruptly or were extended with additional years of mild happiness, they conducted an additional trial asking participants which life they would likely choose, rather than just rating them. They surprisingly found no preference for the shorter life that ended on a high note, despite favouring it in previous ratings. These findings illustrate a crucial distinction between retrospective evaluation and direct choice; the former relies on emotional state, whereas the latter is likely more influenced by immediate reward. Zauberman et al. (2006) reported that humans making more objective, analytical judgments are less affected by an experience’s emotional progression, weakening the peak-end effect. This type of processing may be relevant to nonhuman animals, like bees, where the role of emotional behaviour remains a topic of debate (see Baracchi et al., 2017).

The question of whether bees are susceptible to peak-end effects is also ecologically relevant. In nature, bees often exhibit a bottom-to-top visitation pattern on vertical inflorescences (Heinrich, 1975; Waddington and Heinrich, 1979), giving each visit an “end.” This stereotyped foraging behaviour is thought to help bees avoid costly flower revisits (Pyke, 1979; Best and Bierzychudek, 1982), similarly to traplining in flower patches (Cartar, 2004). If bees respond to peak-end effects, could this influence how nectar is distributed in plants? Floral structures are under selection and probably impacted by pollinator behaviour (Harder et al., 2004; Bailey et al., 2007). Concentrating rewards in top flowers could enhance perceived value while conserving energy, yet this pattern is not common across flowering plants. Conversely, many plants have their older, lower flowers producing higher nectar rewards than their newer, upper flowers (Waddington and Heinrich, 1979; Best and Bierzychudek, 1982). Moreover, studies that reversed the rewards order found that bees maintained their upward visitation pattern, even when top flowers offered the most nectar (Waddington and Heinrich, 1979; Corbet et al., 1981). Bees typically reject and leave flower patches when the rewards decrease (Pyke, 1978; Heinrich, 1979). Specifically, Townsend-Mehler et al. (2011) showed that bumblebees are more likely than honeybees to abandon food sources that are decreasing in value and are quicker to discover new ones. Furthermore, bumblebees have been shown to leave flowers based on the nectar amount in the last flower probed (Cresswell, 1990) and often abandon an inflorescence when nectar levels drop below a certain threshold (Hodges, 1985). Such behaviour may benefit plants by encouraging pollinators to move to other flowers, thereby reducing the risk of self-pollination (Biernaskie et al., 2002). A better understanding of peak-end effects in foraging bees could offer new perspectives on nectar distribution patterns in plants and the co-evolution of plants and pollinators.

## SUPPLEMENTARY MATERIAL

**S1** contains the statistical analysis of the main experiment. **S2** provides the reflectance curves of the artificial flowers, and **S3** includes video clips of bees foraging. The dataset for the experiment is provided in **S4**. Full statistical analyses and datasets for all experiments are also available on Zenodo (https://doi.org/10.5281/zenodo.14068426).

## ACKNOWLEDGMENTS

We want to thank A. Koch, A. Vethacke and J. Burger for their assistance with data collection, and I. Prem for support with video analysis. Special thanks to A. Avarguès-Weber for providing spectrometer measurements of the artificial flowers.

